# Multi-contact 3C data reveal that the human genome is largely unentangled

**DOI:** 10.1101/2020.03.03.975425

**Authors:** Filipe Tavares-Cadete, Davood Norouzi, Bastiaan Dekker, Yu Liu, Job Dekker

## Abstract

The genome is organized into chromosome territories that are themselves spatially segregated in A and B compartments. The extent to which interacting compartment domains and chromosomes are topologically entangled is not known. We show that detection of series of co-occurring chromatin interactions using multi-contact 3C (MC-3C) reveals insights into the topological entanglement of compartment domains and territories. We find that series of co-occurring interactions and their order represent interaction percolation paths through nuclear space in single cells where fragment 1 interacts with fragment 2, which in turn interacts with fragment 3 and so on. Analysis of paths that cross two chromosome territories revealed very little mixing of chromatin from the two chromosomes. Similarly, paths that cross compartment domains show that loci from interacting domains do not mix. Polymer simulations show that such paths are consistent with chromosomes and compartment domains behaving as topologically closed polymers that are not catenated with one another. Simulations show that even low levels of random strand passage, e.g. through topoisomerase II activity, would result in entanglements and mixing of loci of different chromosomes and compartment domains with concomitant changes in interaction paths inconsistent with MC-3C data. Our results show that cells maintain a largely unentangled state of chromosomes and compartment domains.

## INTRODUCTION

As cells exit mitosis the nucleus reforms and individual chromosomes decondense while maintaining their territorial organization ^1^. Within each territory chromosomes become spatially compartmentalized: domains of active and inactive chromatin cluster together to form euchromatic A and heterochromatic B compartments, respectively. Interactions between chromosomes also develop where individual chromosome territories interact and where some level of apparent intermingling can be observed ^2^. As for intra-chromosomal associations, inter-chromosomal contacts are enriched for interactions between domains of the same type (A-A and B-B interactions).

In the crowded chromatin environment of the nucleus any locally acting topoisomerase II enzymes could randomly pass strands through each other, which would lead to topological entanglements between chromatin from different chromosomes or from different domains within chromosomes producing catenations. Given sufficient time, this could lead to complete mixing of chromosomes reaching an equilibrium state ^3, 4^. Considerable theoretical analyses have explored how chromosomes can maintain individual and largely separate territories. In a simulation study by Rosa et al ^4^, segregation and unmixing of chromosomal domains were associated with two generic polymer effects: nuclear confinement (i.e. chromatin density) and topological disentanglement. It was suggested that the topologically disentangled state is the result of slow kinetics of chromatin relaxation after metaphase ^4^. The assumption was that the interphase nuclei is not equilibrated and behaves like a semi-dilute solution of unentangled ring polymers which are known to segregate due to topological constraints ^3–7^. The enormous length of mammalian chromosomes makes the reptation time, i.e. the time for the ends of the linear chromosome to explore the nucleus, extremely long so that chromosomes, and chromosomal sub-domains, can effectively be treated as though they are ring polymers.

Despite the fact that nucleus-wide equilibration and mixing of chromosomes is prohibitively slow, localized mingling, and Topoisomerase II – mediated interlinking between chromosomes and sub-chromosomal domains could still occur. Imaging experiments have shown that neighboring chromosome territories overlap to some extent at their borders and at these locations chromatin from different chromosomes appears to mix. However, it is not known whether this mingling is the result of formation of local entanglements due to topoisomerase II activity. Similarly, active and inactive compartment domains that can be located far apart along chromosomes cluster together to form euchromatic and heterochromatic compartments. Interactions between such distal domains are readily detected by chromosome conformation capture based methods ^8, 9^, but whether this involves or leads to topological entanglements is not known.

Topological entanglements of chromatin fibers could create complications for any process acting on the genome including transcription, replication and chromosome condensation and segregation. However, the extent to which chromosome entanglements form during interphase is not known because experimentally detecting such topological features of chromosomes has not been technically possible. Here we show that detection of strings of co-occurring chromatin interactions in single cells using Multi-Contact 3C (MC-3C) can reveal the extent to which catenations between chromosomes and between chromosomal domains occur. Comparing experimental MC-3C data with polymer simulations we find that chromosomes and chromosomal compartments are largely devoid of entanglements.

## RESULTS

We implemented an experimental method and corresponding polymer simulations to detect and analyze sets of multi-contact chromatin interactions that occur in single cells. The method is based on conventional chromosome conformation capture (3C) ^10^ and the C-walk and multi-contact 4C approaches developed earlier ^11–13^. 3C performed with the restriction enzyme DpnII produces long ligation products where multiple restriction fragments become ligated together to form DNA molecules that can be >6 kb long, and that are mostly linear (Supplemental Figure 1). Such long linear strings of ligation products (referred to as C-walks ^12^) can represent clusters of loci that interact simultaneously with each other in single cells (“clusters”, Figure 1A). C-walks can also represent connected paths of pairwise interactions where fragment 1 interacts with fragment 2, which in turn interacts with fragment 3 while fragments 1 and 3 do not directly interact (Figure 1A). Such connectivity can occur in highly connected interaction networks where each pair of loci can be linked through any number of indirect interactions, referred to as interaction or bond percolation ^14^. Here we will treat C-walks as “percolation paths” to reflect the fact that they can, and often do (see below) represent series of linked fragments.

**Figure 1:**
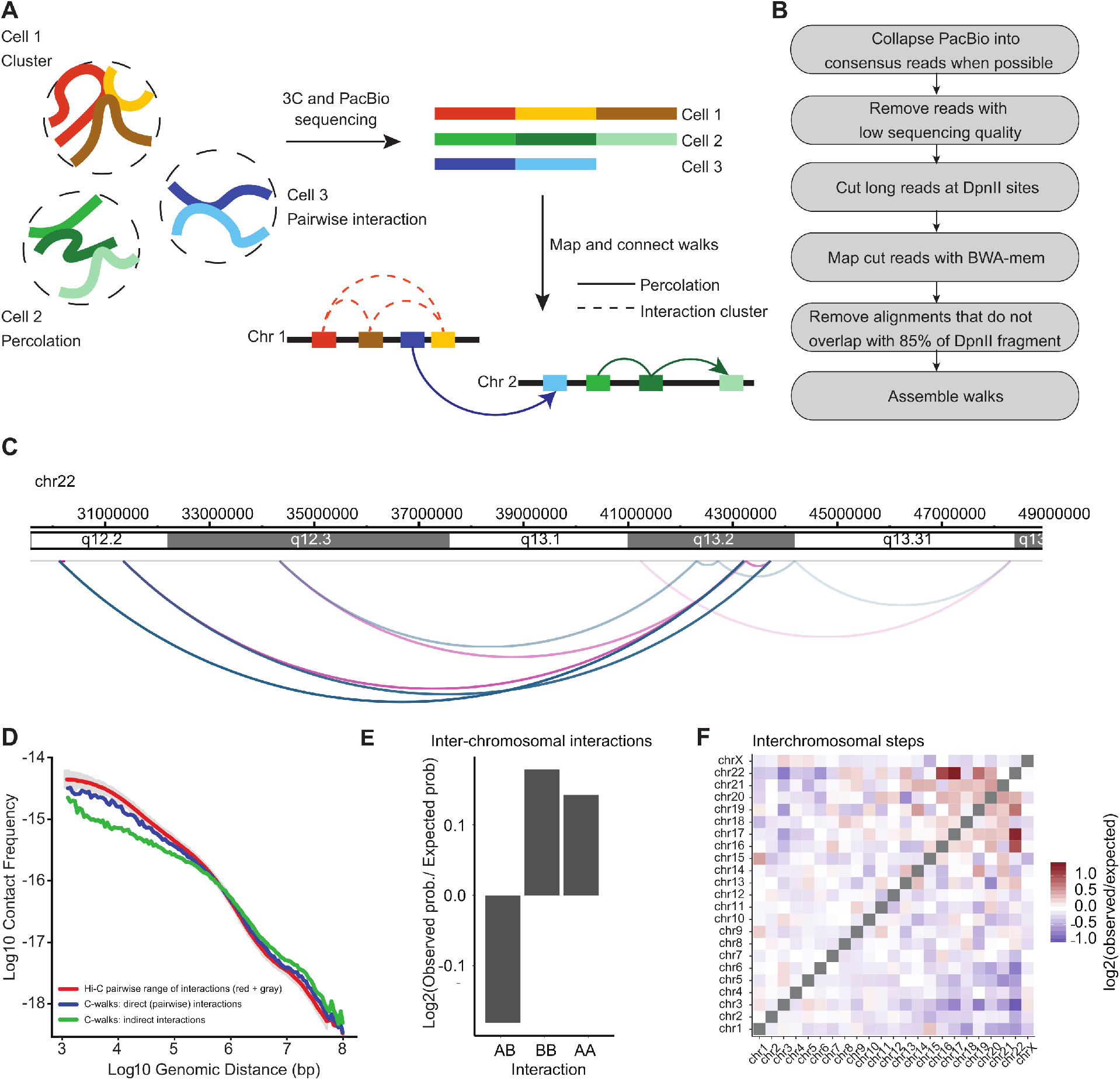
Outline of MC-3C. **A**: Schematic of the MC-3C approach. By performing PacBio sequencing on the products of a 3C experiment fragments that belong to the same interaction cluster or percolation path are captured as chimeric long reads. **B:** Computational pipeline used to process PacBio reading reads to generate C-walks. **C:** Example of a C-walk. Red arcs represent steps that in the positive direction and blue arcs represent steps in the negative direction. The steps get darker during the walk. **D**: Interaction frequency as a function of genomic distance. **E**. Comparison of inter-chromosomal A-A, B-B and A-B interactions. An expected probability of A-A, B-B and A-B interactions was calculated from all detected fragments that participate in inter-chromosomal interactions. This was compared to the observed proportion of interactions and converted to a base-2 logarithm. **F**. Observed/expected number of inter-chromosomal steps. Small chromosomes preferentially interact.

To identify the restriction fragments that make up each string of ligation products DNA generated with 3C was sequenced on a Pacific Biosciences RS II DNA sequencer (Figure 1B). To improve the quality of sequencing error-prone PacBio reads we used the SMRT Analysis v 2.2.0 package (PacBiosciences) to obtain consensus sequences when the same molecule was sequenced more than once and to remove reads with low quality (Figure 1B). Sequence reads were split in individual restriction fragments by computationally splitting reads at DpnII sites. Individual sections were then mapped to the human genome sequence (Hg19) using BWA-mem (^15^, Figure 1B). Two adjacent fragments within one C-walk that map to two adjacent restriction fragments in the genome were merged and treated as a single fragment as they are likely partial digestion products. We further excluded reads that map to <85% of a restriction fragment or that visit the same fragment more than once. Fragments that could not be uniquely mapped were kept as steps in the C-walk. The result is a set of C-walks, or percolation paths, similar to the one depicted in Figure 1C. Each walk is an ordered set of interactions – here named steps – that connect fragments that are located on the same chromosome or on different chromosomes. The physical dimensions of such paths within the cell nucleus are probably in the range of up to several hundred nm: a C-walk where fragments interact end-to-end and combined is 6 kb would maximally be around several hundred nm long (assuming a contour length of the chromatin fiber of 40 nm/kb ^16^) but will likely occupy a volume with a smaller diameter.

We applied MC-3C to exponentially growing HeLa S3 cells that are mostly in interphase. After processing and pooling data obtained from 2 independent biological replicates, we obtained a set of 118,154 interphase C-walks. To verify the quality of the data we first treated the interactions as pairwise data including only direct interactions that are between adjacent fragments within the C-walks. We plotted their interaction frequency (*P*) as a function of genomic distance (*s*, in bp) between the loci (Figure 1D). *P*(*s*) for C-walk interactions is similar to that obtained from a conventional Hi-C dataset down-sampled to the same number of interactions. Interestingly, *P*(*s*) for indirect interactions within C-walks, where all pairwise combinations of fragments within a C-walk are considered an interaction, deviates from the *P*(*s*) of direct interactions in C-walks and pairwise Hi-C data (Figure 1D). There is generally a shift to longer-range interactions. This observation indicates that indirect interactions are not equivalent to direct interactions and therefore that the order of fragments in C-walks is not random. This implies that C-walks are not reflecting sets of chromatin segments that all interact with each other simultaneously as in the cluster model.

### Interaction interfaces between chromosomes are relatively smooth

We first explored the subset of C-walks that include interactions between two different chromosomes, as these sets of co-occurring interactions could provide insights into the structure of the interface where two chromosome territories touch. Similar to previous analyses of C-walks ^12^, and to be conservative, we removed C-walks that include interactions between more than 2 chromosomes because some of these could be the result of random ligation events (48% of inter-chromosomal C-walks). The remaining inter-chromosomal C-walks (42,851 in total) reproduce known features of inter-chromosomal interactions such as enrichment of interactions between A and between B compartments (Figure 1E) and of inter-chromosomal interactions between small, gene-rich chromosomes, as previously described in Hi-C experiments ^9^ (Figure 1F).

We were interested to determine whether the string of ligation products forming inter-chromosomal C-walks would provide information about the nature and structure of the interaction interface of two chromosome territories. If at the interface of two territories loci from each chromosome freely mix, we expect to see multiple inter-chromosomal steps within each C-walk, and loci from the two chromosomes could potentially be ligated in any order (Figure 2A). Such mixing could involve various degrees of interlinking and catenation of the two chromosomes. In contrast when the two territories are proximal but do not locally mix or interlink, e.g. there is a smooth interface, we would expect far fewer steps between the two chromosomes per inter-chromosomal C-walk, while observing many intra-chromosomal steps connecting loci within each of the territories (Figure 2B).

**Figure 2:**
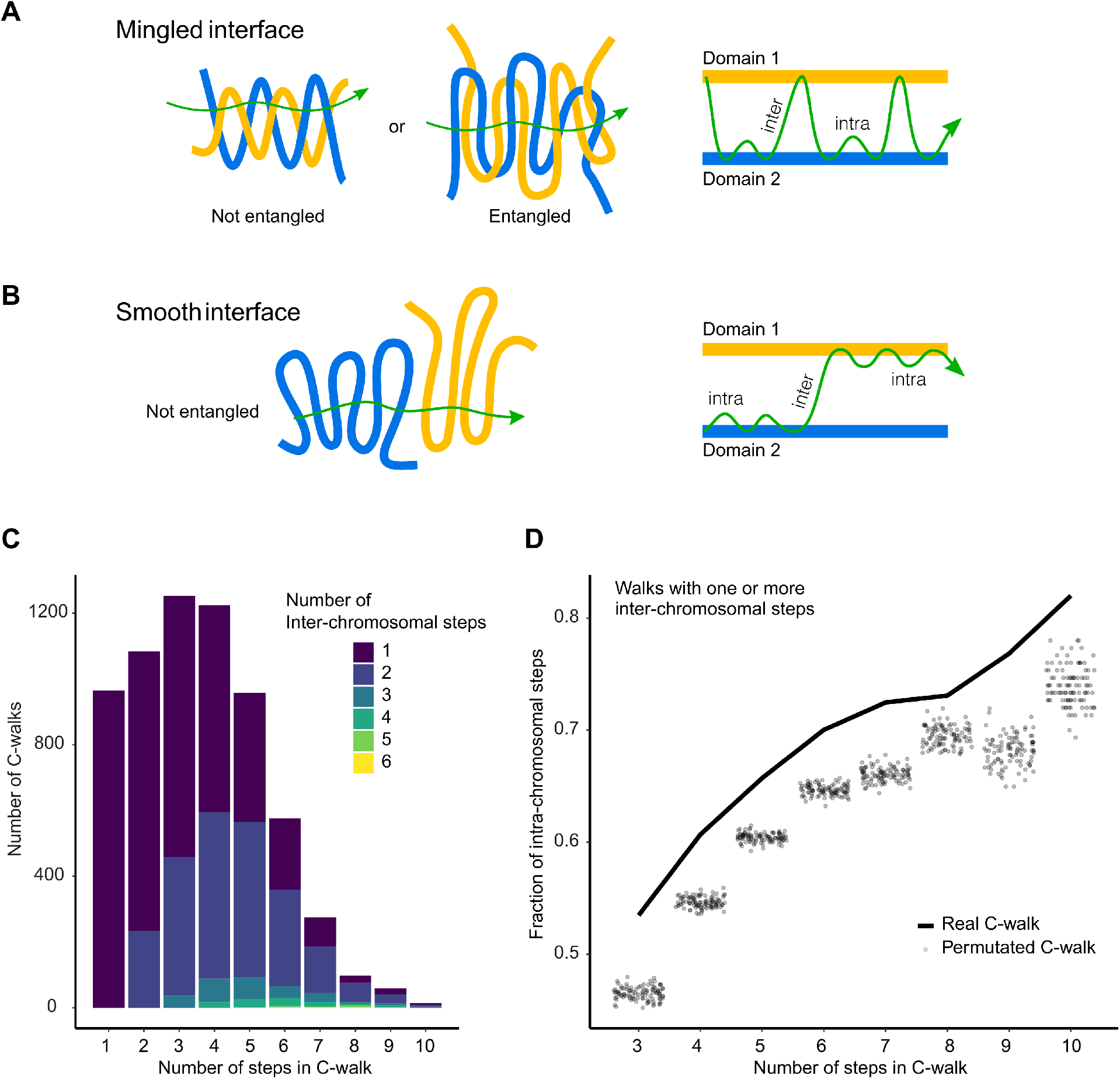
Inter-chromosomal interfaces show limited mixing. **A:** Schematic of how mingled and smooth interfaces would be captured in multi-contact data. In a mingled interface (top row) a C-walk would contain many steps between the two chromosomes, as schematically indicated on the right. In a smooth interface the C-walk would contain series of steps within one chromosomes and only occasionally contain a step between the two chromosomes. **B:** Stacked barplots with the number of C-walks containing at least one inter-chromosomal step separated by the number of steps between two chromosomes (in colors) and the number of steps (along the x-axis) that they contain. Most inter-chromosomal C-walks contain only one or two inter-chromosomal step. **C:** Proportion of intra-chromosomal steps in the set of C-walks with one or more inter-chromosomal steps, separated by the total number of steps in each C-walk. The solid line shows the proportion for real C-walks. The dots correspond to the same proportion in 100 sets of permutated walks. The observed C-walks have a higher proportion of intra-chromosomal steps than would expected if the order of the steps was random.

We performed two analyses to distinguish between these two possibilities. First, we determined the number of inter-chromosomal steps that occur within each inter-chromosomal C-walk (Figure 2C). Interestingly, most inter-chromosomal C-walks only have 1 or 2 inter-chromosomal steps, even for long C-walks, with the remaining steps before or after this step occurring between fragments contained within either territory. C-walks of increasing length tend to add mostly intra-chromosomal steps, and as a result the percentage of intra-chromosomal steps per C-walk increases as compared to shorter C-walks (Figure 2D). Second, we explored whether the order of the fragments within each C-walk mattered. For this analysis we only used C-walks for which all fragments could be mapped. We created 100 sets of permutations (for each set of C-walks of length *n* (number of steps)) in which we randomized the order of DpnII fragments within observed inter-chromosomal C-walks, and then for each permutated set again calculated the percentage of intra-chromosomal steps per C-walk. We find that real C-walks have more intra-chromosomal steps than permutated walks, for C-walks of all lengths (Figure 2D). The same phenomenon is observed when C-walks were analyzed separately for inter-chromosomal interfaces between two A domains, two B domains or between an A and a B domain (Supplemental Figure 2A). Importantly, one implication of this result is that the order of fragments in C-walk ligation strings is meaningful and that they do not simply represent clusters of fragments that all interact with each other at the same time. Instead, many C-walks likely represent percolation paths.

The explanation for the observation that randomizing the order of steps within inter-chromosomal C-walks leads to lower numbers of intra-chromosomal steps is that in real C-walks steps within each chromosome are clustered together to form continuous intra-chromosomal sub-walks, and the C-walk only infrequently crosses from one chromosome territory to another. Randomization of the order of steps within C-walks breaks these continuous sub-walks within each chromosome and mixes fragments from the two chromosomes so that more steps will be between fragments located on different chromosomes. These results indicate that at the interface of two chromosome territories there is limited mixing of chromatin fibers from the two chromosomes and the interface between the two territories is rather smooth, at least at the scale of a few hundred nanometers. We further note that the span of the intra-chromosomal interactions (i.e. the largest intra-chromosomal distance between two fragments in the C-walk ^12^) is bimodal, with enrichments of distances in the range of hundreds of kb and in the range of several Mb (Supplemental Figure 2B). This distribution is similar to the overall distribution of all intra-chromosomal steps and represents interactions within and between compartment domains respectively (see below). This indicates that, near the point of contact between chromosomes, loci separated by large intra-chromosomal distances and located in different compartment domains can co-localize.

### Within chromosome territories compartment domains interact but do not mingle

Next, we explored properties of C-walks that occur within chromosome territories. We first determined the step size distribution. We find a bimodal distribution, with enrichments of steps in the range of hundreds of kb, and in the range of several Mb (Figure 3A, gray line). Such distribution has been observed before ^12^. We find that the shorter-range steps involve interactions between fragments located within a single compartment domain (either A or B; Figure 3A, blue line), while the longer-range steps involve interactions between loci located in different compartment domains (either A-A, B-B, or A-B interactions; Figure 3A, red line). 43.4% of all intra-chromosomal steps involve interactions between fragments located in different compartmental domains, and 87% of all intra-chromosomal C-walks involve more than one compartmental domain, indicating extensive contact between distal compartment domains. As expected, we find that interactions between two A compartment domains or two B compartment domains occur more frequently than interactions between an A and a B compartment (Figure 3B).

**Figure 3:**
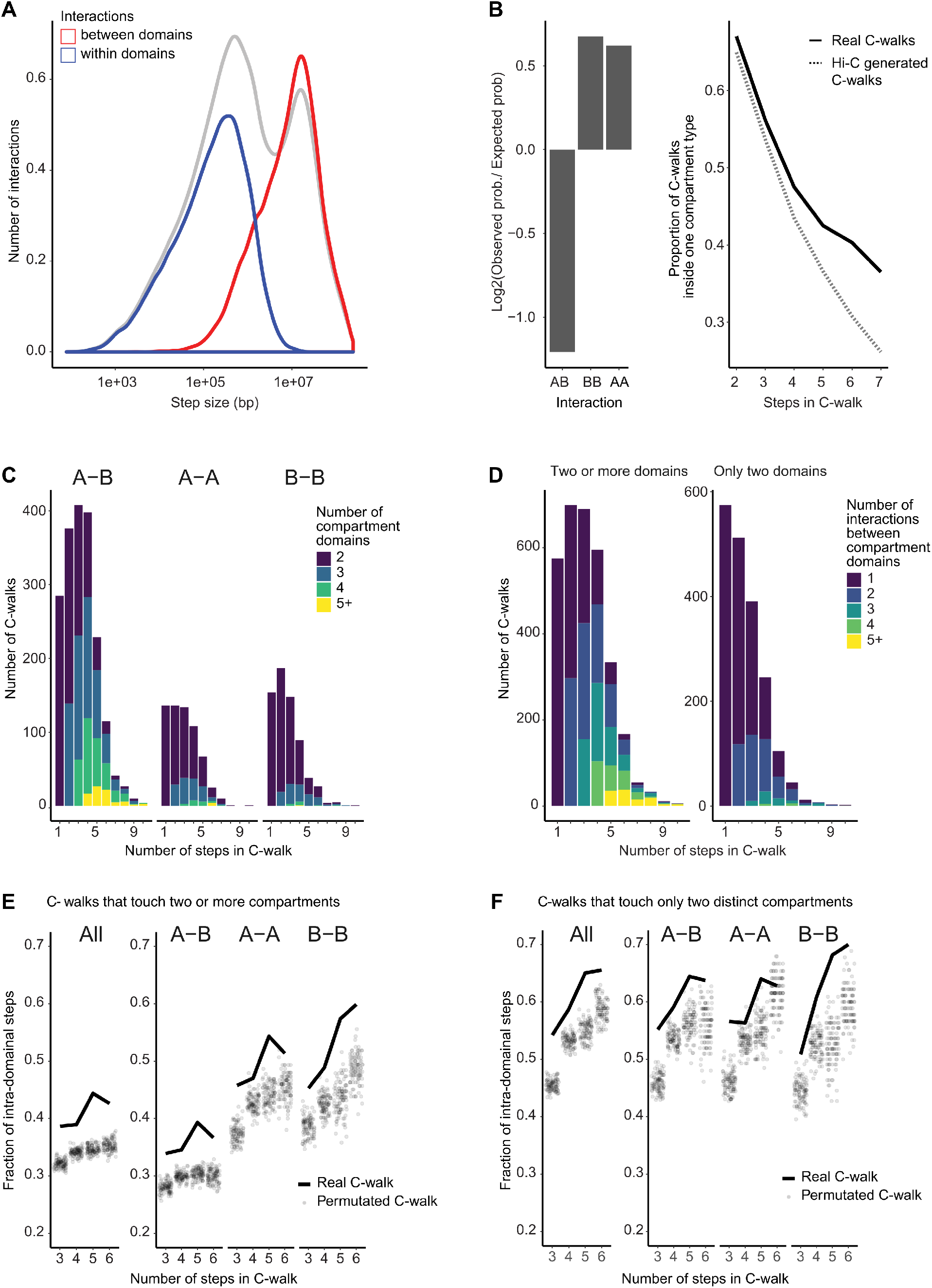
Intra-chromosomal interfaces between compartment domains show limited mixing. **A:** Density plot of the distance of all (gray line), intra-domain (blue line) and inter-domain interactions (red line, including A-A, B-B and A-B interactions). The area under the gray curve of all interactions has been adjusted so that it is double of the area under the curve of the each of the red and blue lines. Only interactions with valid distances (intra-chromosomal) have been used. **B:** Left panel: Comparison of A-A, B-B and A-B interactions. An expected probability of A-A, B-B and A-B interactions was calculated from all detected fragments that participate in intra-chromosomal interactions. This was compared to the observed proportion of interactions and converted to a base-2 logarithm. Right panel: For each number of steps, the proportion of C-walks that only contact domains of one compartment type (A or B) was calculated. This was done for real multi-contact data (full line) and for a set of computationally generated C-walks that was produced from sampling pairwise Hi-C data. These were done by choosing one 1kb bin at random from chromosome 4. Then one among the bins it interacts with was randomly selected, weighted by the number of interactions. This becomes the first step of the simulated C-walk. Then from this new bin another interaction was selected in the same manner, and so on. This was done for the same number of C-walks and steps as the original C-walk data. **C:** Stacked barplots with the number of compartment domains traversed by intra-chromosomal C-walks that visit 2 of more domains. These are separated by the number of steps in the C-walk on the x-axis and further separated into subplots what show C-walks that traverse both A and B domains (A-B), only A domains (A-A) and only B domains (B-B). C-walks that visit five or more domains were grouped together in the plotting. **D:** Stacked barplots showing the number of intra-chromosomal C-walks that visit exactly two distinct domains (right subplot) and for intra-chromosomal walks that visit two or more distinct domains (left subplot) with 1, 2, 3, 4 and five or more inter-domain steps, separated on the x-axis by the total number of steps in a C-walk. **E:** Proportion of intra-domain steps in C-walks that visit two or more domains, separated on the x-axis by the total number of steps in each C-walk (solid line). These are compared to the same calculation done on 100 sets of permutated walks (semi-opaque dots). Each of one hundred permutated set of walks was done by taking real C-walks and reordering the interactions in each walk, while keeping the same fragments. This was done for all the C-walks that touch two or more domains (on the left), and separately for those that visit both A and B domains (A-B), those that visit only A domains (A-A) and those that visit only B domains (B-B). **F:** Same as panel E but using the set of C-walks (and corresponding permutations) that visit only two compartment domains.

We computationally generated a comparable set of intra-chromosomal walks by sampling from pairwise Hi-C data. Starting with a single step, steps were added by random sampling from the set of pairwise Hi-C interactions of the last added fragment in the walk (see Methods). We find that for any given number of steps C-walks are more likely to stay within the same compartment type – visiting just A or just B compartment territories – than the Hi-C derived walks (Figure 3B, right panel). This again indicates that steps in C-walks are inter-dependent, consistent with previous comparisons between multi-contact data and pairwise data ^12^.

C-walks that involve interactions between two compartmental domains likely represent sets of interactions at the interface where two or more of these domains touch. Such C-walks can provide insights into the structural features of these interfaces, i.e. the extent to which distal compartment domains mingle where they interact. We first determined the number of distinct compartment domains that are captured within the subset of intra-chromosomal C-walks that involve interactions between at least 2 distinct compartment domains. We find that such C-walks can involve interactions between up to as many as 11 distinct compartment domains dependent on the number of steps within the C-walk (Figure 3C). Such C-walks often are between sets of either A or B domains, as would be expected, but with one or a few steps in a compartment domain of opposite type mixed in. We noticed that such domains of opposite type tend to be located directly adjacent in the linear genome to one of the other compartment domains of concordant type that are part of the same C-walk. This indicates these are likely not random ligation events. When we analyze C-walks that exclusively involve only A or only B compartment domains, we find that such inter-compartment C-walks typically contain fragments from up to 4 distinct domains (Figure 3C).

To test whether the order of fragments in intra-chromosomal C-walks matters we again employed permutations. We randomized the order of fragments within C-walks that involve at least two compartment domains along the same chromosome (irrespective of compartment type, Figure 3D left plot) and calculated the number of steps that occur within individual compartment domains (Figure 3E). For this analysis we again included only C-walks for which all fragments could be mapped. We find that real C-walks have more intra-compartment domain steps than randomized C-walks. This was found for C-walks that include fragments from both A and B compartments, only A compartments or only B compartments (Figure 3E). Similar results were found when we restricted our analysis to C-walks that involve only two compartment domains (A-A, B-B and A-B interactions; Figure 3F). This result shows again that the order of steps within real C-walks is meaningful. The fact that real C-walks display more intra-compartment domain steps suggests limited intermingling of chromatin at the interface of two interacting compartment domains so that for each locus the nearest neighbor tends to be located within the same compartment domain, similar to what we observed for chromosomal interfaces.

### Polymer simulations show that MC-3C data are consistent with unentangled chromosome and domain interfaces

The limited mixing of chromatin from different chromosomes or sub-chromosomal domains could reflect a lack of topological entanglement. Topological entanglement would increase the mixing at the interface ^3, 17^. To test this hypothesis, we performed coarse-grained simulations of chromosome and domain interaction interfaces with and without topological entanglements and then determined how topological differences at the interface affect chromatin mixing and the composition of C-walks.

We first simulated chromatin domains as topologically closed polymers (Supplemental Figure S3). In these simulations the ends of each polymer, each representing a chromosomal domain located on the same or on different chromosomes, were held together making them effectively rings. For sufficiently long polymers, local sub-chains can be treated as topologically closed systems ^18^. Weak attractions between monomers were included to simulate the attractive forces leading the A and B compartment formation ^19, 20^, although including these did not affect the results (Supplemental Figure S4). We also tested the effect of chromatin density (from dilute to crowded) but found that this did not affect the results either (Supplemental Figure S4). Simulations included sufficient numbers of Monte Carlo steps to reach equilibrium (Supplemental Figure 3C). Figure 4A shows the interface between two topologically closed polymers in yellow and cyan, reflecting two interacting chromosomal domains that can be located along the same chromosome or on different chromosomes. Unlinking is guaranteed by keeping the domain ends, shown in red and black ball pairs, fixed and proximate. This snapshot shows that after millions of Monte Carlo steps, the two domains remain largely unmixed.

**Figure 4:**
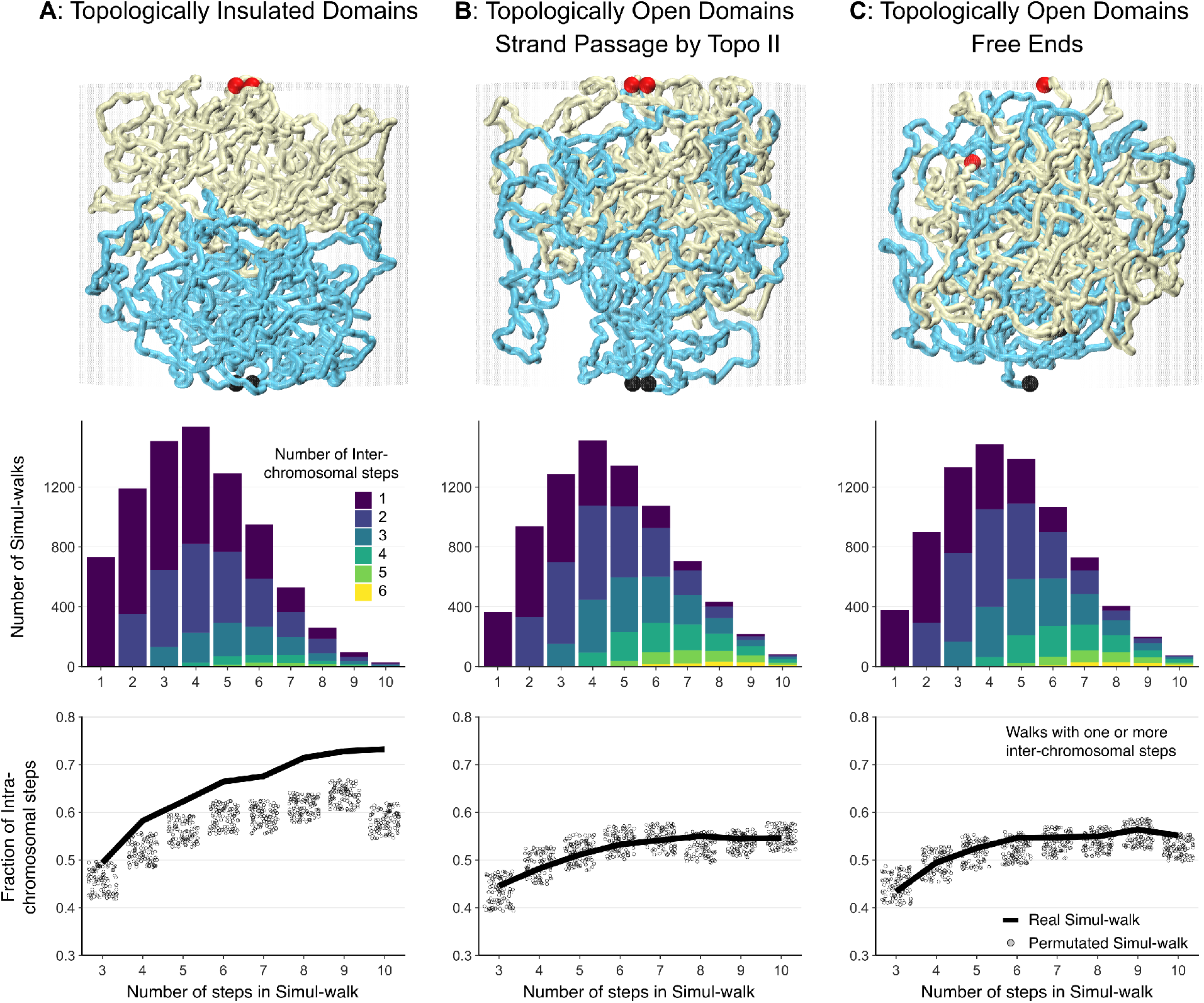
Impact of topological linking and entanglement on mixing at compartment domains or chromosomal interfaces. Simulations were performed on three types of interfaces: (**A**) Topologically unlinked domains, (**B**) Topologically linked domains via strand-passage, and (**C**) Topologically open domains via free-end movements. Chains are shown in yellow and cyan with ends highlighted by the red and black balls. The interface remains smooth and unmixed for the topologically insulated domains in panel A, while the two topologically open domains in panels B and C exhibit considerable mingling. The interface mixing of the simulated chains is quantified by generating, computationally, percolation paths similar to the C-walks obtained by MC-3C. These are called simul-walks. In the second row, the number of inter-chomosomal (or inter-domain) steps that occur within each inter-chain simul-walk is presented. In simulation A, most simul-walks involve only 1 or 2 inter-domain interactions similar to the C-walks in Figure 2B. In simulations in panel B and C the number of simul-walks with only 1 inter-domain interaction has dropped while simul-walks with 3 to 6 inter-domain crossings are now much more likely. Histograms in panels B and C (middle) indicate that the two chains are largely mingled. In the bottom row, the fraction of intra-domain steps in simul-walks with at least one inter-domain interaction is shown in solid black lines (compare to Figure 2D and Figure 3E, F for C-walks). The unmixed interfaces in simulation A demonstrate a higher fraction of intra-domain steps in simul-walks with at least one inter-domian crossing. Randomly permutating those same simul-walks results in a lower fraction of intra-domain steps shown by gray dots. In contrast, for the mixed and catenated interfaces in simulations in panels B and C, the fraction of intra-domain steps in simul-walks is indistinguishable from permutated simul-walks indicating extensive mixing.

Next we simulated C-walks as percolation paths through the simulated polymer systems (see Methods). Briefly, we randomly chose a location along either of the two polymers and then stepped with a given step size (*r*_cutoff_ = 75 nm) to a proximal polymer section. We generated a collection of simulated C-walks (simul-walks) in this manner and selected the subset that contained at least one inter-domain step. We then performed the same set of analyses and permutations on this sub-set of inter-domain (or inter-chromosomal) simul-walks as we did for experimental C-walks (Figures 2 and 3). The number of inter-domain steps that occur within inter-domain simul-walks are shown in the middle row in Figure 4. Interestingly, most inter-domain simul-walks contain only 1 or 2 inter-domains steps with the remaining steps before or after this step occurring between fragments within either domain, very similar to what we found in the experimental C-walks between chromosomes and between compartment domains within chromosomes (Figures 2 and 3). When we permutated the steps within this set of simul-walks we observed a reduction in the number of intra-domains steps, corresponding to more inter-domain steps (Figure 4A, bottom). This simulation reproduces very closely what we observed when we permutated experimental C-walks (Figure 2 and Figure 3). The results from the simulations did not change when we used different sizes of the steps (*r*_cutoff_) for calculating simul-walks (Supplemental Figure S5).

When we performed this analysis for polymer systems that were topologically open (Figure 4B and 4C) we obtained very different results. First, in the presence of strand passage or free movement of ends of the interacting domains, we observed extensive mixing and catenation of the two interacting polymers (Figure 4B, 4C, top row; Supplemental Figure 3C) ^21^. Second, simul-walks generated under these conditions displayed features very different from experimental inter-chromosomal or inter-compartment domain C-walks. The number of simul-walks with 1 inter-domain step decreased, while walks with 3, 4, 5 and 6 inter-domain steps increased dramatically (Figure 4B and 4C, middle row). Permutating steps within these simul-walks did not change the number of intra-domain steps. This indicates that intra-domain steps within these simul-walks are not forming clustered sub-walks within each domain, consistent with the observation that the domains are extensively mixed (Figure 4B and 4C, top row). Combination of these analyses indicate that experimental C-walks are consistent with an absence of entanglements between chromosomes or between compartment domains.

### Analysis of C-walks contained within compartment domains

To measure the level of mixing of chromatin within compartment domains we analyzed C-walks that involve interactions between two segments of 250 kb each that are separated by 0.5, 1.0 and 2.5 Mb, and that are both contained within the same compartment domain. This analysis allows investigation of the extent to which two distal chromatin segments within a single compartment domain mix and is analogous to the analysis shown in Figure 3 for interactions between compartment domains. The results are presented in Figure 5. We plotted the fraction of intra-segmental steps as a function of the length of the C-walk and compared this to permutated C-walks, exactly as we did for intra-chromosomal C-walks that connect different compartment domains (above). We observed that the fraction of intra-segmental steps in the majority of permutated C-walks is lower than for experimental C-walks, but that there is overlap as well. This indicates that segments separated by 0.5-1.5 Mb but located within the same compartment domain can display considerable, but possibly not complete, mixing.

**Figure 5:**
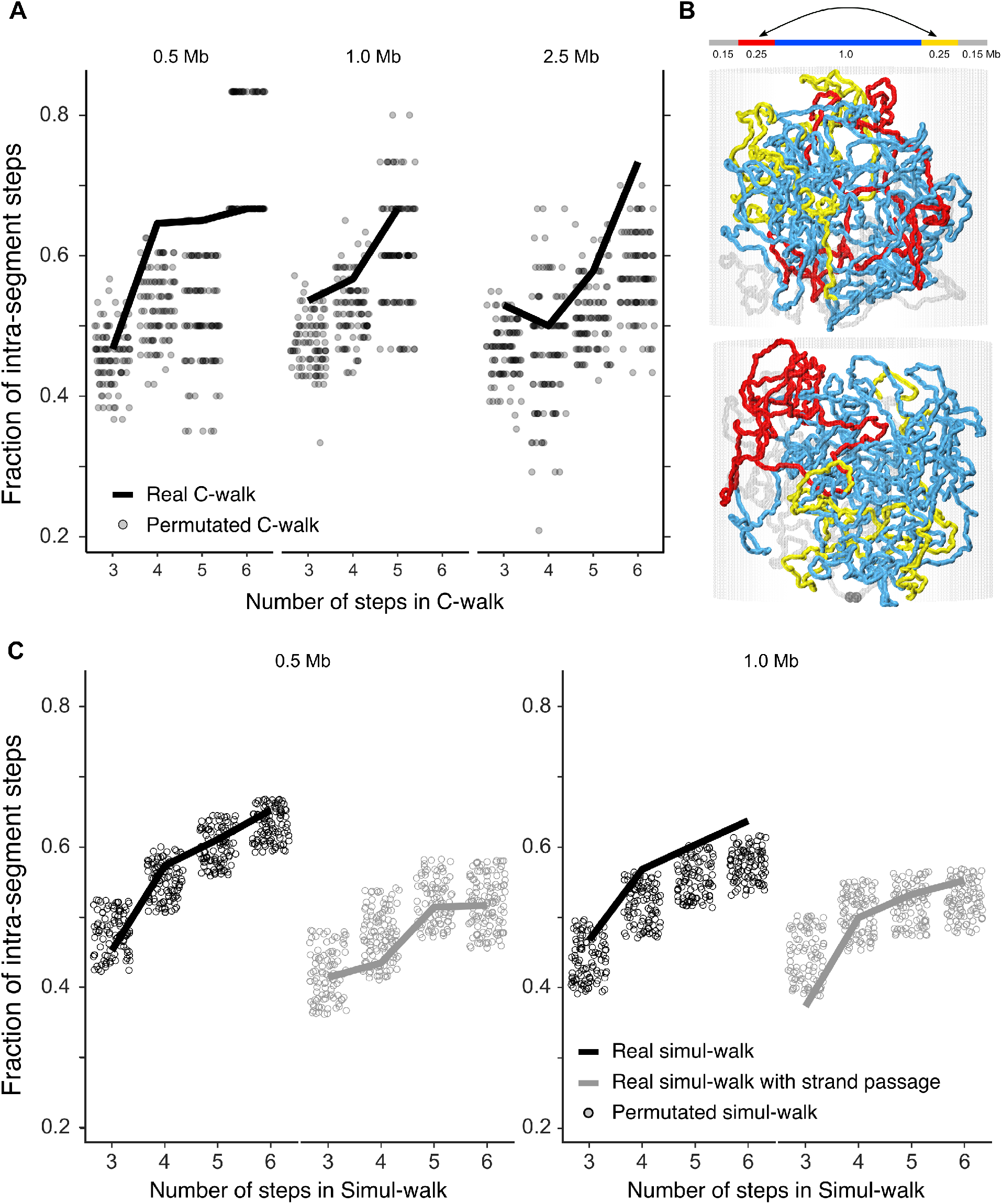
Distal parts within the same compartment domain can show considerable mixing but may also have low levels of strand crossing. **A**. Analysis of C-walks connecting two segments of 250 kb each, spaced by 0.5, 1.0 and 2.5 Mb and that are contained within a single compartment domain. The proportion of intra-segment steps in this set of C-walks is shown, separated by the total number of steps in each C-walk. Very few C-walks had more than 6 steps and are left out. The solid line shows the proportion for real C-walks. The dots correspond to the same proportion in 100 sets of permutated walks. The observed C-walks have a higher proportion of intra-chromosomal steps than the majority permutated sets, but there is overlap. The overlap indicates that the two distal segments are mixed to some extent. **B**. Top panel: schematic depiction of the polymer chain used for simulations in the middle and lower panels. Middle and lower panels: snapshots of the system with the two 250 kb segments highlighted in red and yellow and the middle 1.0 Mb segment in cyan. Snapshots show that inside a domain the chromatin is mixed and that the 250 kb end segments are not long enough to produce blob-like conformations that would make them impenetrable with respect to each other. On the other hand, the 1.0 Mb cyan fragment is sufficiently long and generates a topological blob and barrier for mixing with the rest of the chromatin. **C**. Analysis of simul-walks obtained with polymer simulations depicted in B with and without allowing strand passage. The fraction of intra-segment steps in real simul-walks (solid lines) and permutated simul-walks (circles) are more similar indicating mixing between the fragments. When segments of 250 kb are separated by 1.0 Mb, mixing is reduced when strand passage is not allowed, and the statistics of such simul-walks most resemble experimental C-walks shown in A.

To determine the extent to which mixing is expected to occur at this length scale, and how this depends on allowing strand passage, we performed the following simulations. Compartment domains were simulated as circular molecules within a confined space (semi-dilute regime, Supplemental Figure S4). For sufficiently long polymers sub-chains can be treated as closed rings (see above). We do not assume any active folding mechanisms to act such as loop extrusion in these simulations (see discussion). We performed Monte Carlo simulations with (topologically open) or without (topological closed) strand passage. Figure 5B shows simulation snapshots in the absence of strand passage with the two 250 kb segments highlighted in red and yellow and the middle 1.0 Mb region in cyan. Snapshots show that inside a domain the chromatin is mixed and that the 250 kb segments are not long enough to produce blob-like conformations that would make them impenetrable with respect to each other. On the other hand, the 1.0 Mb cyan region produces a topological blob and barrier for mixing with the rest of the chromatin. These simulations show that chromatin at short length scales (hundred of kb) is a random polymer chain which readily mingles while at larger scales (Mb) chromatin resembles a collection of unmixed neighboring blobs with their surfaces in smooth contact.

We then calculated simul-walks through these polymer conformations as described above, and selected simul-walks that involved at least one step between the two distal segments of 250 kb that are separated by 0.5 or 1.0 Mb (red and yellow domains, separated by cyan segment in Figure 5B). We then calculated the fraction of such intra-segmental steps in these simul-walks and compared that to the fraction of intra-domain steps observed after permutating the simul-walks (Figure 5C). We observe that for domains separated by 0.5 Mb the fraction of intra-segmental steps in simul-walks is comparable to that for permutated simul-walks, both in the absence or presence of strand passage. This indicates that domains that are separated by relatively small distances (0.5 Mb) readily mingle even in the absence of strand passage. We note that in the absence of strand passage the fraction of intra-segmental steps of simul-walks is larger than that observed when strand passage is allowed. Interestingly, a similar larger fraction of intra-segmental steps is observed in experimental C-walks, which may indicate that chromatin within compartment domains display limited knotting or entanglement.

Next, we analyzed simul-walks and permutated simul-walks that involve interactions between segments separated by 1.0 Mb. As expected, in the presence of strand passage the fraction of intra-segmental steps in simul-walks is comparable to that of permutated simul-walks, indicating extensive mixing of the segments. However, in the absence of strand passage, the fraction of intra-segmental steps for permutated simul-walks is in the majority of cases lower than for the simul-walks. This indicates that segments separated by 1.0 Mb can mix to some extent even in the absence of strand passage, but that strand passage leads to more extensive mixing. Comparing the results from the simul-walks to those for experimental C-walks we note that the observations with experimental C-walks (including for segments separated by 0.5 Mb) most resemble those of the simul-walks obtained from simulations for domains separated by 1.0 Mb in the absence of strand passage. This suggests that distal regions within compartment domains mix to some extent but display few if any entanglements.

## DISCUSSION

We present experimental data that show that the genome is largely topologically unentangled. A summary of our findings is depicted in Figure 6. Figure 6A portrays chromatin as an organization of largely unentangled blob-like domains in semi-crowded conditions with rather smooth interfaces. Figure 6B illustrates the same domains with a large degree of entanglement, such that the boundaries between domains and between chromosomes fade away. Experimental inter-chromosomal and inter-compartment domain C-walks are consistent with a lack of topological entanglements between chromosomes and between compartment domains, as illustrated in Figure 6A. At shorter length scale, e.g. within compartment domains and in the absence of active mechanisms, we found that chromatin becomes mixed to some extent even without allowing strand passage and therefore our approach has less power to discern presence or absence of topological transitions. However, experimental C-walks within compartment domains best resemble predicted simul-walks for topologically fixed polymer states even for segments separated by relatively small genomic distances (0.5 Mb). For such small separations our simulations predicted more mixing (Figure 5). Possibly active processes, not included in the simulations, play roles in disentangling chromatin at smaller scales (see below). Overall, our data lead us to propose that the genome is largely unentangled at all length scales.

**Figure 6:**
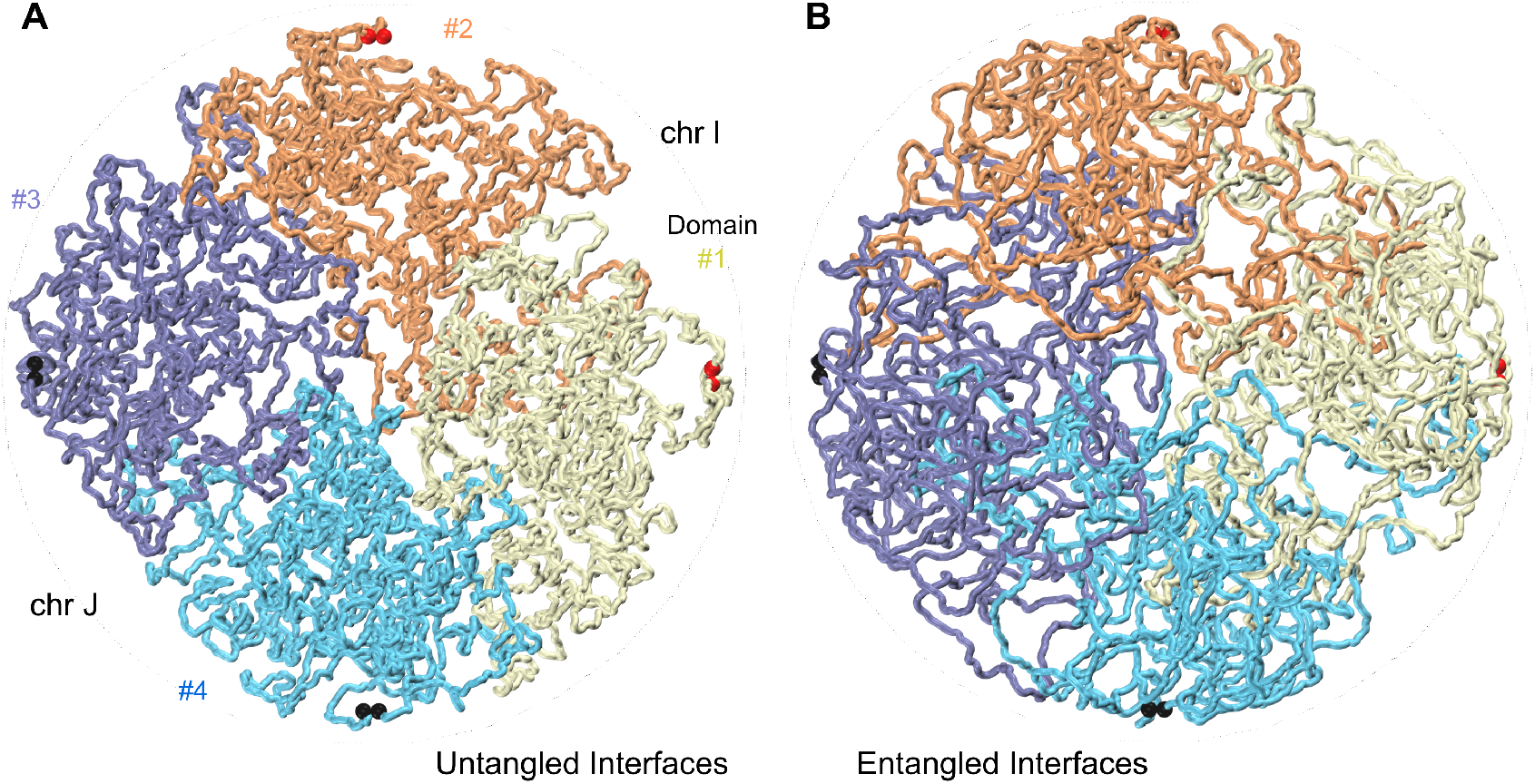
Chromatin is an organization of largely untangled blob-like domains squeezed together under a semi-crowded condition with rather smooth interfaces. **A**. Untangled chromosomal and domain interfaces. Four domains #1 to #4, two pairs from different chromosomes I and J, are pushed together in a cylindrical container. The interfaces are smooth and untangled. Figure B shows the same system with strand passage allowed. Chromatin domains diffuse and fuse into each other and distinct chromosomal and domain interfaces, visible in A, fade away.

Although theoretical considerations and polymer simulations had suggested that chromosomes would display low levels of entanglement in cycling cells for kinetic reasons, it has proven difficult to experimentally assess the topological state of the genome in cells directly. Previous studies employed multi-contact data to explore the presence of sub-nuclear compartments ^22^, and to test whether interactions between genes and regulatory elements occur in hubs or in pairs ^11–13^. Here we find that the overall statistics of the composition and order of multi-contact data can also provide insights into the topological nature of the interfaces between chromosomes and chromosomal domains revealing a global lack of chromosomal entanglements. Similarity of the statistics between experimental C-walks and the simul-walks shows that the two major factors responsible for the smooth interfaces between domains and chromosomes are nuclear crowding (confinement) and topological disentanglement. However, unlike previous reports ^4^, we observe these phenomena even under well-equilibrated conditions for the typical domain sizes in contact. Similar to previous reports, the unmixing applies to a wide range of domain sizes and no sign of a crossover to any mixed regime is observed ^7^. Lack of entanglement within contiguous domains was recently reported by Goundaroulis and co-workers using an entirely independent approach: their re-analysis of oligo-paint serial fluorescent labeling data from Bintu and colleagues ^23^ indicates chromatin domains can be largely knot free, at least for the relatively small subset of cells where the chromatin fiber could be reliably traced ^24^. This study did not investigate the extent to which different chromosomes or different chromosome compartments domains are entangled.

MC-3C data can reveal entanglements at chromosome and domain interfaces because it allows distinguishing patterns of intra-chromosomal interactions occurring at the interfaces between chromosomes or sub-chromosomal domains from those occurring farther from these interfaces, which is not possible with pairwise 3C-based data. The reason MC-3C allows this distinction is that intra-chromosomal interactions occurring near such interfaces tend to be contained within C-walks that also contain at least one inter-domain or inter-chromosomal step. MC-3C data show that intra-chromosomal and intra-domain interactions near interfaces occur in clusters that are relatively rarely broken up by an inter-chromosomal or inter-domain interaction, consistent with limited mixing. Our MC-3C data also show that the order of steps within C-walks is meaningful, and that strings of interactions represent percolations paths through crowded nuclei. This is a critical consideration when using such data to study interaction hubs, e.g. between genes and multiple distal regulatory elements.

Previous imaging studies found that chromosome territories overlap at least partially ^2, 25^. Combined with our data these observations indicate that chromatin fibers, or chromosomal domains from one chromosome can locally invade another territory to some extent, but apparently without becoming topologically linked and entangled. The overlap between chromosome territories observed microscopically occurs at a length-scale of several microns, while percolation paths studied here are probably in the range of hundreds of nanometers. It will be interesting to acquire much longer C-walks that cover up to microns in 3D space to determine whether any additional architectural features of territory boundaries can be detected. This will require isolating much longer 3C ligation products, combined with very long read sequencing platforms such as nanopore sequencing.

In a crowded environment of the nucleus, and with topoisomerase II acting locally to randomly pass strands, chromosomes would form a highly entangled and knotted state, at least at their interfaces ^26^. Our data strongly support a very different state in which chromosomes and compartment domains are not entangled. How is entanglement prevented when topoisomerase II is abundantly present? One possibility is that topoisomerase II is acting only rarely throughout the genome. Given that in mitosis the genome is largely disentangled, reformation of the interphase genome conformation in the absence of strand passage would then prevent entanglements from forming. Some entanglements can occur through reptation of the ends of the chromosomes, but when chromosomes are very long, as in humans, this effect would be limited ^4^. However, there is evidence that Topoisomerase II is acting throughout the genome. Topoisomerase II-mediated breaks can readily be detected at CTCF sites and gene promoters ^27, 28^. Thus, it appears other processes must act to ensure that entanglements are prevented or are selectively removed, possibly by making topoisomerase II act not randomly so that it is directed towards disentanglement. Recent theoretical analyses have demonstrated that a process of loop formation through chromatin extrusion that strictly acts *in cis* will lead to a largely disentangled genome ^29–31^. Polymer simulations have shown that formation of arrays of loops during mitosis will drive segregation and decatenation of sister chromatids ^32, 33^. Similar extrusion processes in interphase are predicted to position linkages and crossings between and along chromosomes such that topoisomerase II action will remove these links leading to topological simplification ^30, 31^.

Cohesin complexes have been proposed to extrude loops in vivo during interphase, while condensin complexes generate dense arrays of extruded loops in mitosis. The process of loop extrusion in interphase was first proposed based on patterns of Hi-C data representing topologically associating domains (TADs) and detection of loops between convergent CTCF sites ^34–38^, while loop extrusion has been a logical mechanism for generating arrays of loops observed along mitotic chromosomes ^39–43^. Recent in vitro experiments have demonstrated that cohesin and condensin complexes indeed can extrude loops in an ATP-dependent mechanism ^44–47^. These observations provide strong support for the model in which loop extrusion events throughout the cell cycle guide topoisomerase II activity to generate and maintain a largely decatenated genome as we detect here. MC-3C combined with polymer simulations can provide a powerful experimental approach to assess the topological state of chromosomes, e.g. as cells progress through the cell cycle, and in cases where cells display a variety of chromosome folding defects.

## ACKNOWLEDGMENTS

We thank members of the Dekker and Mirny labs for helpful discussions. We acknowledge support from the National Institutes of Health Common Fund 4D Nucleome Program (DK107980), and the National Human Genome Research Institute (HG003143). J.D is an investigator of the Howard Hughes Medical Institute.

## METHODS

### Cell Culture

Hela S3 cells were cultured in Dulbecco’s Modified Eagle Medium (DMEM, Gibco, 10569044) supplemented with 10% Fetal Bovine Serum (Gibco, 16000), 100 U/ml penicillin (Gibco, 15140) and 100 μg/ml streptomycin (Gibco, 15140) at 37°C and 5% g/ml streptomycin (Gibco, 15140) at 37°C and 5% CO2.

### Multi-contact 3C protocol

In situ Chromosome conformation capture (3C) was performed as previously described ^10^, with modifications described in Belaghzal et al. ^48^. Briefly, cells were washed with HBSS (Gibco, 14025092), cross-linked with 1% formaldehyde (Fisher, BP531) for 10 minutes at room temperature. Crosslinking was quenched by addition of glycine to a final concentration of 125 mM and cells were incubated for 5 minutes at room temperature, followed by incubation on ice for 15 minutes. Aliquots of 5 million cross-linked cells were lysed by incubation in lysis buffer (10 mM Tris-HCl (pH=8.0), 10 mM NaCl, 0.2% Igepal CA-630) supplemented with Halt protease inhibitor (Thermo Fisher, 78429) for 15 minutes on ice. Cells were disrupted with a dounce homogenizer using pestle A (2x 30 strokes). A final concentration of 0.1% SDS was added and extracts were incubated at 65°C for 10 minutes. SDS was quenched by addition of with Triton X-100 to a final concentration of 1%. Chromatin was then digested with 400 U DpnII (NEB, R0543) at 37°C overnight. After enzyme inactivation by incubation at 65°C for 20 minutes, DNA ligation was performed by addition of 10 µL T4 DNA ligase (NEB, M0202) and incubation at 16°C for 4 hours. Crosslinking was reversed by addition of 50 µl proteinase K (10mg/ml) (Invitrogen, 25530031) followed by incubation at 65°C for 2 hours, followed by another addition of 50 µL proteinase K (10mg/ml) and overnight incubation at 65°C. DNA was isolated by 1:1 phenol/chloroform (Fisher, BP1750I) extraction followed by ethanol precipitation. RNA was removed by addition of and 1 µL RNase (1 mg/ml; Sigma, 10109169001) and incubation at 37°C for 30 minutes. Prior to sequencing on Pacbio RS II the 3C library was size-selected with Bluepippin.

### Determination of topology of 3C ligation products

Pacbio sequencing relies on adapter ligation and therefore any circular ligation products in 3C libraries could not be sequenced. To assess whether strings of 3C ligation products are linear or circular we treated 3C libraries with exonuclease V (NEB, M0345) or T5 exonuclease (NEB, M0363). These nucleases only degrade linear DNA. As a control plasmid pCMV6 plasmid (4.6kb) was linearized by digestion with NdeI (NEB, RO111). 180 ng control linearized plasmid DNA and 180 ng of 3C library was treated with either 0.5 Unit exonuclease V (NEB, M0345) or 0.5 Unit T5 exonuclease (NEB, M0363) at 37°C for 30 minutes. Degradation of DNA was then analyzed by running samples on a 0.8% Agarose gel. Both exonucleases degraded the linearized plasmid and the 3C ligation product library indicating 3C products are linear. Circular plasmid DNA was not degraded (Supplemental Figure S1).

### PacBio library preparation

Samples for PacBio library preparation were size selected into two groups of small (3-6 kb) and large (6 kb) molecules. Libraries were constructed using the PB Express 2.0 Kit according to the manufacturer’s instructions and sequenced in a PacBio RSII instrument.

### Data processing

#### Extract consensus reads of insert and quality filter

To extract the reads of inserts from the raw PacBio data files we used smrt tools (SMRT Analysis v 2.2.0 package, PacBiosciences). When possible, these correspond to the consensus sequence of several read-throughs of the same DNA molecule. Only reads of insert with a quality threshold over 80 were selected.

#### Virtually digest reads at GATC sites

The extracted reads of insert were virtually cut at the sequence sites recognized by DpnII (GATC) using BioPython scripts.

#### Map reads with BWA-MEM

Cut reads were mapped using bwa-mem ^15^, with default parameters. Pairs of reads that are consecutive on the PacBio molecule and that align with 85% of a genomic DpnII fragment are counted as a direct interaction, or a step. When two adjacent fragments map to two adjacent DpnII fragments in the genome they were merged as these may represent partial digestion products. The different steps that come from a single PacBio molecule constitute a walk. If there was an unmapped cut read in the middle of a walk, a NA was inserted. Walks that visit the same DpnII genomic fragment more than once were filtered out. For downstream analysis two sets of walks were used: those with NA fragments and those without.

#### C-Walks generated from Hi-C data

C-walks were computationally generated from non-synchronous HeLa S3 pairwise Hi-C data. The Hi-C experiment was performed as described ^48, 49^. For each computationally generated walk, one 1 kb bin from chromosome 4 was selected at random. Then, a new fragment was randomly selected from the set of cis interactions of that bin in the Hi-C dataset, excluding the immediate neighbors. From this second fragment the same selection was done, and so on until walks with the desired number of steps were generated. Computationally generated walks were made so that they had the same number of step distribution as the experimental C-walks.

#### Permutated C-walks

Permutated walks were generated by taking all the fragments involved in a C-walk and randomly rearranging the order of the fragments in the walk. One hundred permutations were done per walk. Only C-walks with no NAs were used.

#### Contact Probability for C-walks and down-sampled Hi-C data

To calculate the contact frequency (*P*) as a function of genomic distance *s* we used pair of interacting loci mapped to autosomal chromosomes only. As comparison, HeLa S3 Hi-C data was down-sampled for autosomal chromosomes only for 200 times to obtain sets of equal number of interactions as number of directed by MC-3C (Direct interactions: 101,521; indirect interactions: 219,966). Interactions were selected for genomic distances starting at 1 kb up to 100 Mb using log-binning. The observed number of interactions in each genomic distance bin was divided by total number of possible interactions for in each genomic distance bin.

#### Compartment calls

A-B-compartmentalization profile was calculated by principle component analysis of Hi-C dataset obtained from non-synchronized HeLa S3 cultures binned at 250 kb resolution. The first eigenvector represents the A-B-compartmentalization.

### Downstream analysis and code availability

The C-walk assembly pipeline and scripts necessary to generate all plots are available on https://github.com/dekkerlab/MC-3C_scripts

### Polymer simulations and generation of simul-walks

To better understand the geometry and mixing at the interface of two interacting compartment domains (within or between chromosomes), we simulated the interaction of two domains of length 1.5 Mb using polymer modeling. Polymers were represented as a chain of monomers with harmonic bonds and a bending persistence length of 40 nm, a repulsive excluded volume potential, and an additional small short-range repulsion/attraction (representative of good versus bad solvent conditions) between and across both domains. We typically simulated two 3,000 monomer chains, with one monomer corresponding to 0.5 kb (∼3 nucleosomes), with the width of 30 nm and the bead to bead distances corresponding to an average nucleosome density of 2.5 nucleosomes per 11 nm ^16^. The main simulator is written in Python with the intensive potential function calculations being done through a FORTRAN to Python interface wrapper. The details of coarse-graining, energy terms, Monte Carlo moves, and equilibration tests are given in Supplemental Figure S3.

Three types of domain-domain interfaces were simulated. (1) Topologically insulated domains with fixed-ends and no strand-passage allowed. (2) Topologically open domains with fixed-ends but with strand-passage allowed. This mimicks the activity of the Topoisomerase II enzyme, and (3) topologically open domains with no strand-passage but with freely moving ends. All simulations were started from a non-mingled state with the two domains being pushed towards each other along a cylindrical axis of symmetry.

For every condition, 200 million Monte Carlo steps were performed. From the final 150 million equilibrated steps, 1,000 uncorrelated snapshots of the system were used for statistical averaging and analysis of interaction simul-walks. The results shown, in the main figures, correspond to semi-dilute conditions with a chromatin-chromatin self-interaction energy of E_attraction_ = – 0.05 k_B_T/bead. This weak self-attraction (or bad solvent condition) can give rise to compartmentalization ^20^. Using good solvent conditions along with the different levels of crowding produced similar interface mixing behavior as shown in Supplemental Figure S4.

To generate an ensemble of multi-contact walks from the simulations, referred to as simul-walks, the following steps were performed: <u>Step 1</u>: from a random snapshot of the Monte Carlo ensemble choose one random position on a random chain. <u>Step 2</u>: from the spatially close neighbors to this point, within a cut off distance (*r*_cutoff_) and NOT from the nearest neighbors along the same chain, choose one interaction partner. <u>Step 3</u>: continue step 2 until no new neighbor is available within *r*_cutoff_ to interact with. This is considered one simul-walk. <u>Step 4</u>: Repeat from step 1. From this simul-walk ensemble, histogram of the inter-chromosomal or inter-domain steps and the fraction of intra-chromosomal or intra-domain steps in the walks with at least one inter-chromosomal or inter-domain crossing, are calculated. We checked that the statistics of simul-walks are independent of the parameters such as *r*_cutoff_ (Supplemental Figure S5).

