## Supplemental Materials for "Multi-contact 3C data reveal that the human genome is largely unentangled"

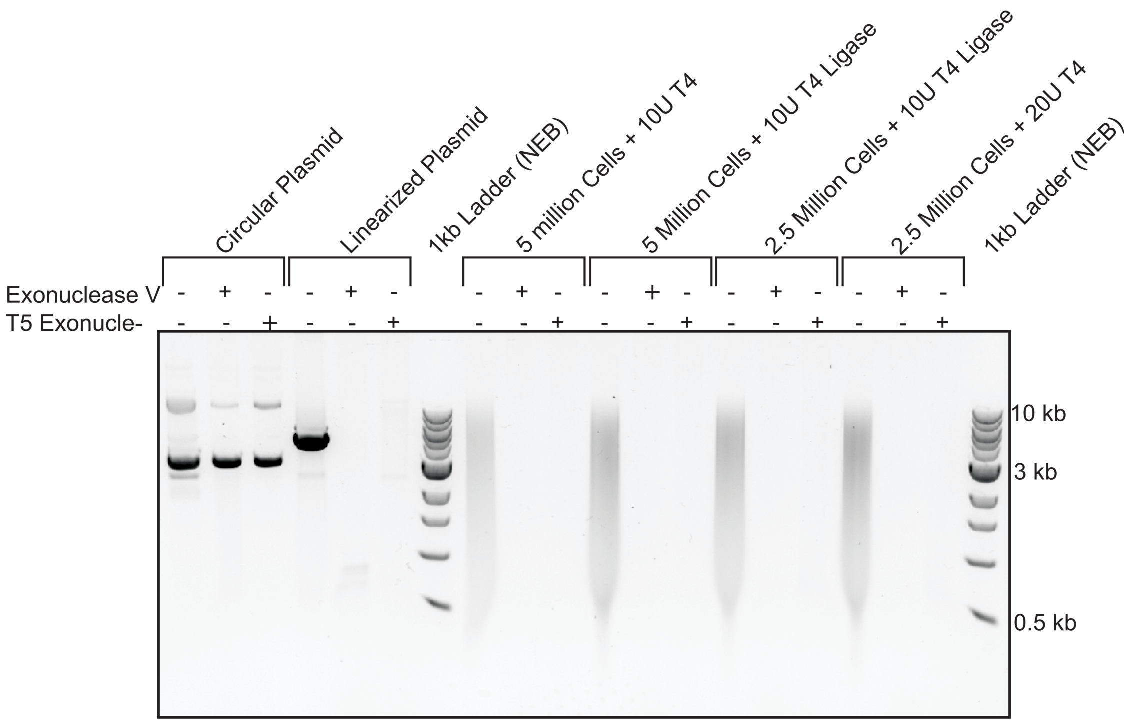


**Supplemental Figure S1**: Agarose gel analysis (0.8% agarose) of DNA size of control circular plasmid DNA, control linear plasmid DNA and of 3C library DNA before and after exonuclease digestion. 3C library and control plasmid DNA were digested with Exonuclease V and T5 exonuclease to determine whether 3C ligation products are linear or circular. In the presence of either exonuclease dsDNA will be digested, and circular DNA will remain present. As control both exonucleases are shown not to digest circular plasmid (pCMV6, 4.6kb) (lane 1-3), whereas linearized plasmid was digested (lane 4-6). Different concentration of 3C library and ligase did not result in detection of circular ligation products as no DNA was detected after exonuclease treatment (lanes 7-18). Markers: 1 kb ladder (New England Biolabs).

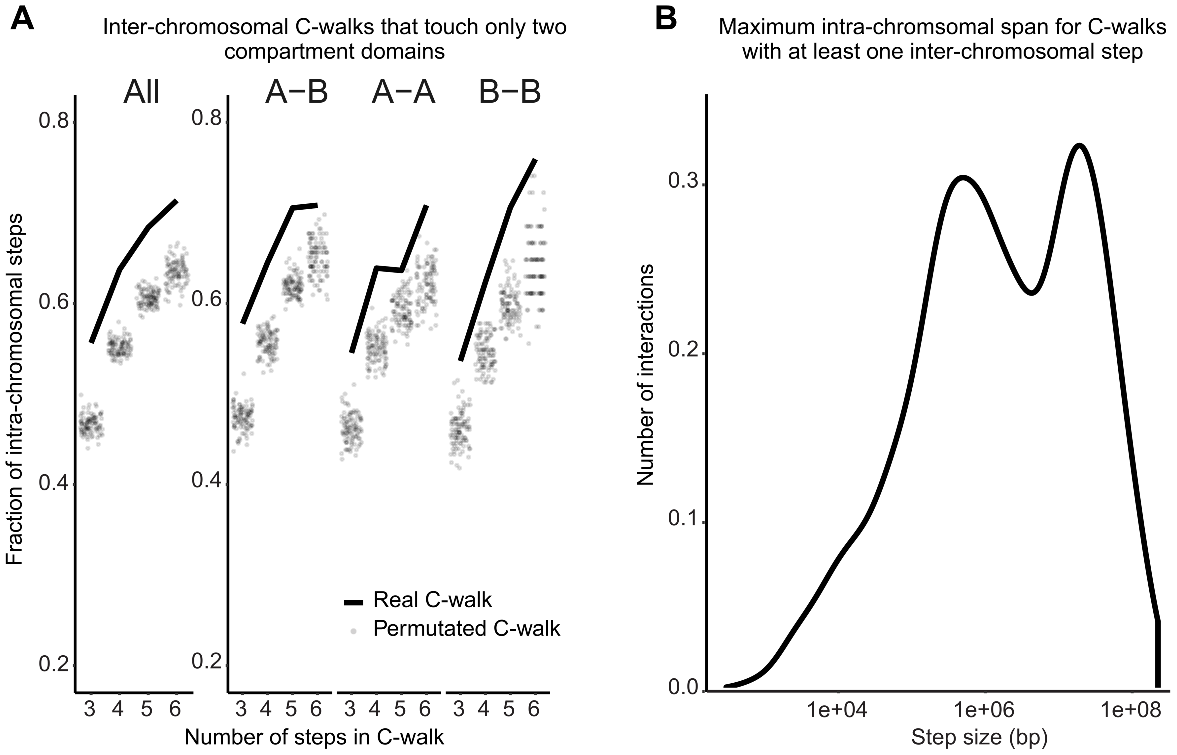


**Supplemental Figure S2**: Additional analyses of inter-chromosonal C-walks. **A**. Proportion of inter-chromosomal steps for C- walks that visit only two distinct compartment domains, separated on the x-axis by the total number of steps in each C-walk (solid line). These are compared to the inter-chromosomal steps after permutating this set of C-walks (dots). This was done for all the walks that touch two domains (on the left), and separately for those that visit one A and one B domain (A-B), those that visit two A domains (A-A) and those that visit two B domains (B-B). **B**. Density plot of the maximum intra-chromosomal span (largest intra-chromosomal distance between fragments that make up a C-walk) for inter-chromosomal C-walks.

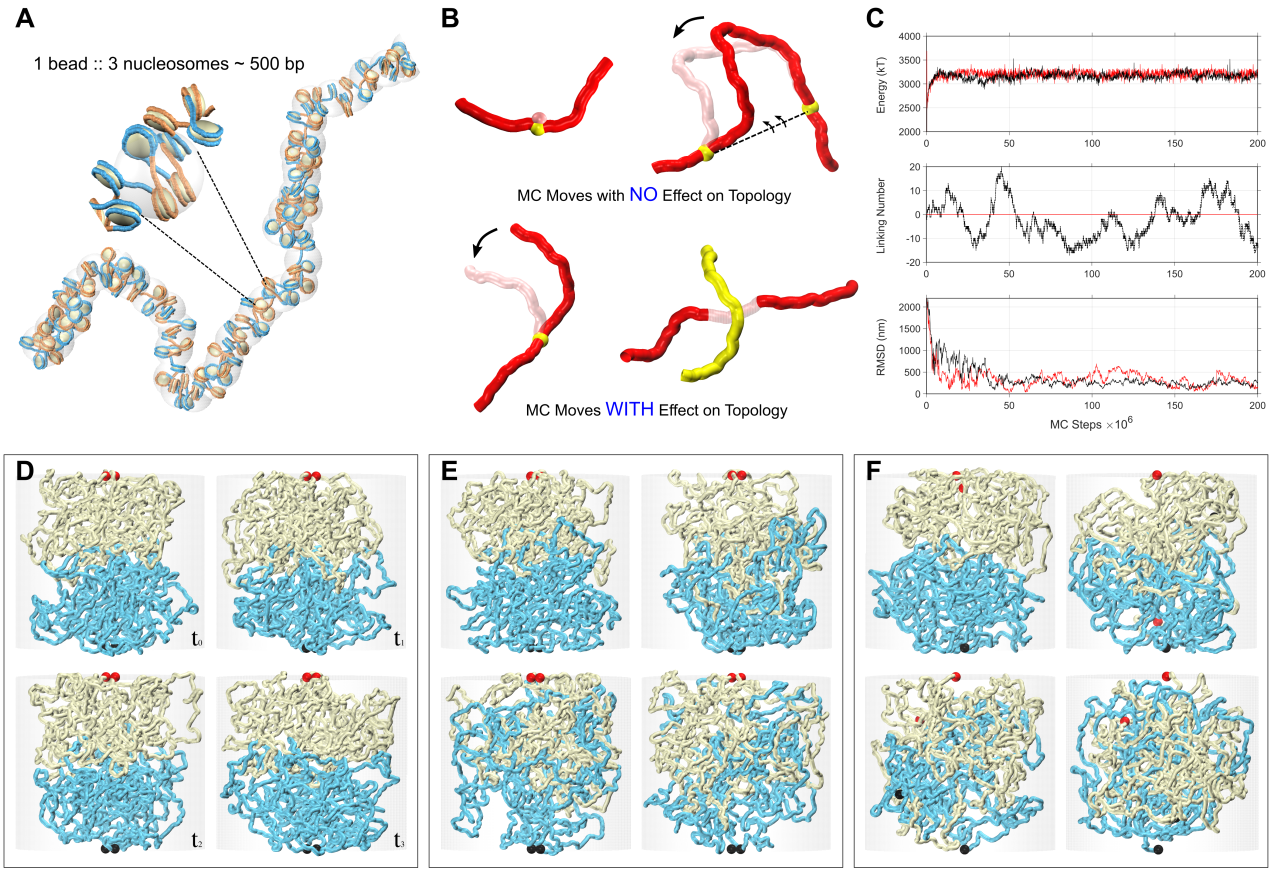


**Supplemental Figure S3**: Details of the Monte-Carlo approach used to study the interface of interacting chromosomal domains in contact. **A**. Coarse-graining is done at the level of chromatin fibers with the diameter of ~25 – 30 nm. Every ~3 nucleosomes are considered one bead with the bead to bead bond distance of ~15 nm at equilibrium and the bending persistence length of 40 nm [50]. A hard-core excluded volume and a short-range pairwise interaction with the form $E_{pairwise} = q \times \exp[-2(r - \sigma)^2 / \sigma^2)]$ is considered, in which, $\sigma$ is the diameter of the beads and "q" is a small negative/positive charge depending on the attraction/repulsion between the beads. **B**. Monte Carlo is used because it is much more effective in sampling the phase space of possible conformations. Two types of Monte Carlo moves are possible. Those which don't change the topology (i.e. the linking or knotting state) of the system and those which do.

Top two moves, i.e. a single bead random movement and a crank-shaft rotation between two randomly chosen points along the chain, do not cause strand passage and do not move the relative position of the chain ends, hence do not affect the topology. Random rotations around a single point and/or brief random ghosting (no excluded volume) of parts of the chromatin fiber allow for relative movement of the ends and/or strand-passage, and thereby affect the topology. **C**. Equilibration for the case with topologically closed domains is presented in red. Equilibration for the case with topologically open domains with fixed-ends but with strand-passage is presented in black. Energy and linking number approach equilibrium after a few millions Monte Carlo steps. Linking number or number of entanglements is always an integer and remains zero for the case with topologically closed domains. RMSD values measured between two distal parts of a chain are shown. It takes up to ~50 million steps for the system to relax in a dynamic-globule structure. We call this a dynamic-globule because even at equilibrium, fluctuations in the distance between two regions on the same chain remain considerable (≈±100 nm). **D**. Simulation snapshots for the case of topologically closed domains with fixed-ends and no strand-passage in “Monte Carlo” time from $t_0$ to $t_3$. The interface between the two domains remains unmixed. **E**. Simulation snapshots for the case of topologically open domains with fixed-ends but with strand-passage allowed. This mimicks the activity of the Topoisomerase II enzyme. Mixing increases gradually as a function of the Monte Carlo time. Note that the domain ends, highlighted with red and black ball pairs, do not move. **F**. Simulation snapshots for the case of topologically open domains with no strand-passage but with freely moving ends. Domains mingle gradually as a function of the Monte Carlo time.

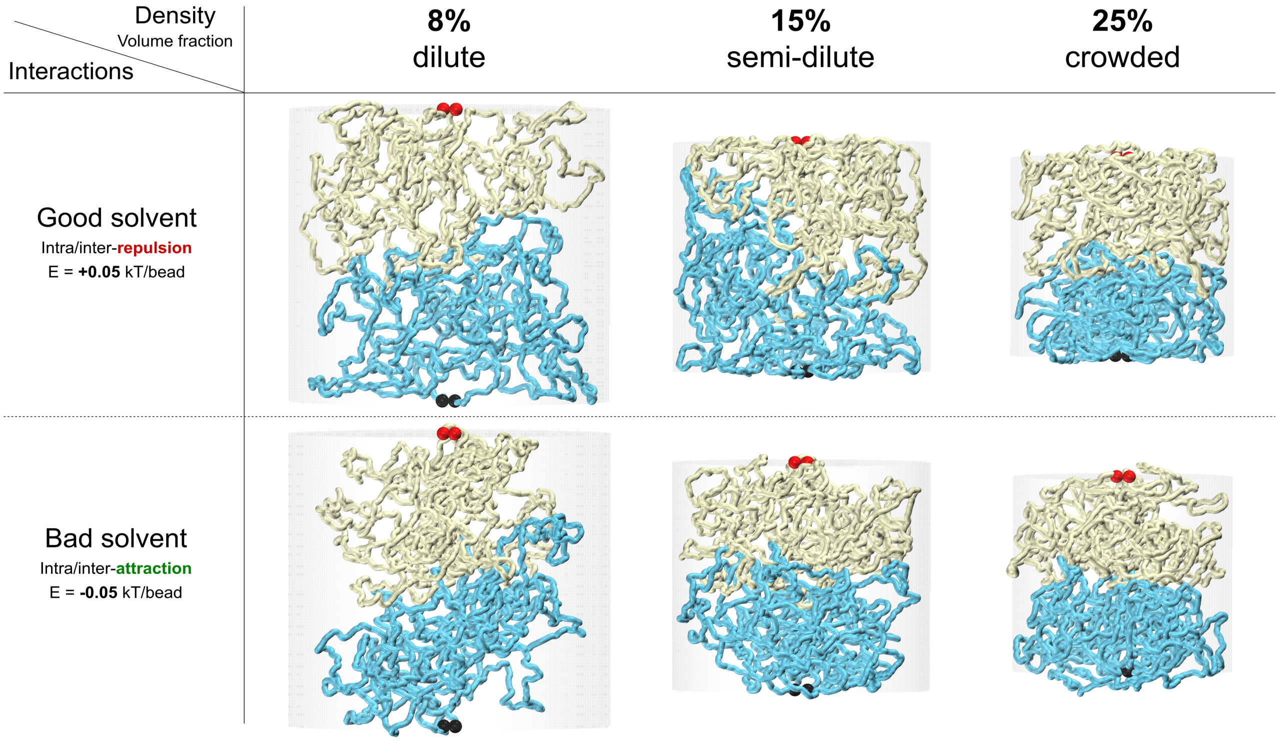


**Supplemental Figure S4**: Equilibrated simulation snapshots for domain interfaces at different solution conditions. Two major determinants of polymer mixing are (**1**) whether polymer prefers to interact with the solvent (good solvent) or to interact with itself (bad solvent). This can be simulated effectively by a repulsive/attractive short-range interaction between the chain's beads. Both conditions might be present in the nucleus leading to microphase separation of chromatin into euchromatin versus heterochromatin domains; and (**2**) the density of the polymer mixture, here simulated by the confining cylindrical walls surrounding the two chains. Our simulations at different solvent and under different crowding conditions still show that domains that are topologically unlinked will barely mix and retain their smooth interfaces. The result might be different for the good solvent in ultra-dilute condition, in which, domains will barely touch and won't generate measurable multi-contact walks. We believe this condition is not relevant to the chromatin *in vivo* and would not be consistent with our MC-3C data.

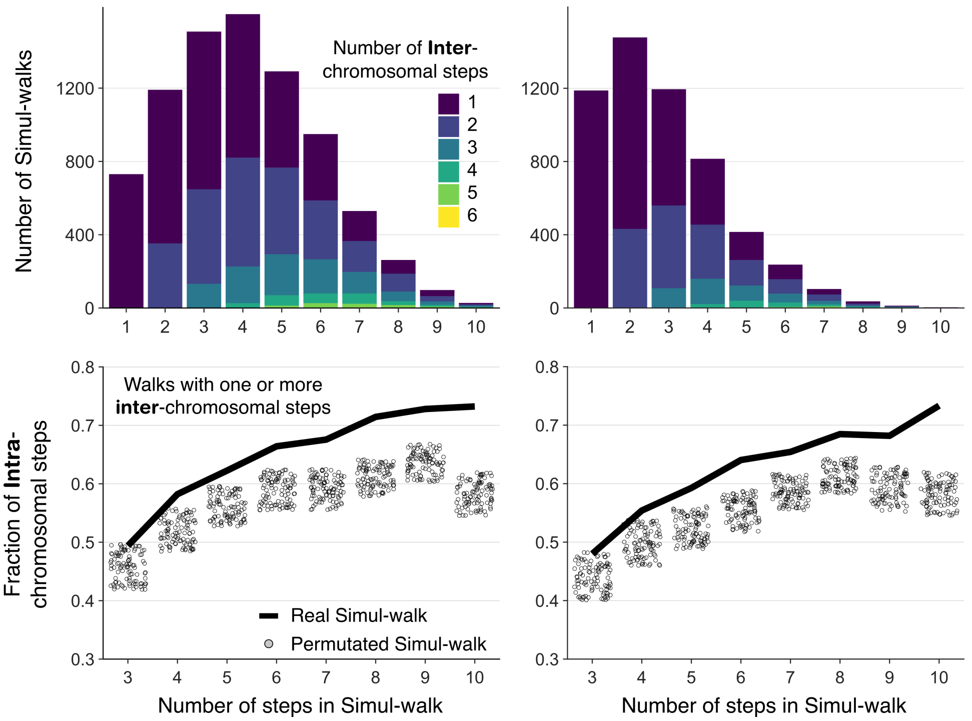


**Supplemental Figure S5**: Simul-walks statistics do not change considerably by changing the selected percolation parameter $r_{cutoff}$ (Left: $r_{cutoff}$ = 75 nm, Right: $r_{cutoff}$ = 60 nm). The left panel is shown in the main Figure 4 because the $r_{cutoff}$ is chosen in such a way that the maximum number of steps in simul-walks in the histogram is ~3-4 which is similar to the peak in the histogram of C-walks in Figure 2C. Regardless, the top two histograms show that most walks still involve only 1 or 2 inter-chromosomal or inter-domain interactions. At the bottom, the shift between the simul-walks (black solid lines) and randomly permutated versions (circles) remains the same as well. The $r_{cutoff}$ value cannot be changed much more than this, because the minimum center to center distance of two touching beads is ~50-60 nm and choosing larger cutoffs will result in skipping some neighboring chains in the percolation path search. Therefore, our results are independent of the parameters we used to find the simul-walks.
